# Timing of host feeding drives rhythms in parasite replication

**DOI:** 10.1101/229674

**Authors:** Kimberley F. Prior, Daan R. van der Veen, Aidan J. O’Donnell, Katherine Cumnock, David Schneider, Arnab Pain, Amit Subudhi, Abhinay Ramaprasad, Samuel S.C. Rund, Nicholas J. Savill, Sarah E. Reece

## Abstract

Circadian rhythms enable organisms to synchronise the processes underpinning survival and reproduction to anticipate daily changes in the external environment. Recent work shows that daily (circadian) rhythms also enable parasites to maximise fitness in the context of ecological interactions with their hosts. Because parasite rhythms matter for their fitness, understanding how they are regulated could lead to innovative ways to reduce the severity and spread of diseases. Here, we examine how host circadian rhythms influence rhythms in the asexual replication of malaria parasites. Asexual replication is responsible for the severity of malaria and fuels transmission of the disease, yet, how parasite rhythms are driven remains a mystery. We perturbed feeding rhythms of hosts by 12 hours (i.e. diurnal feeding in nocturnal mice) to desynchronise the host’s peripheral oscillators from the central, light-entrained oscillator in the brain and their rhythmic outputs. We demonstrate that the rhythms of rodent malaria parasites in day-fed hosts become inverted relative to the rhythms of parasites in night-fed hosts. Our results reveal that the host’s peripheral rhythms (associated with the timing of feeding and metabolism), but not rhythms driven by the central, light-entrained circadian oscillator in the brain, determine the timing (phase) of parasite rhythms. Further investigation reveals that parasite rhythms correlate closely with blood glucose rhythms. In addition, we show that parasite rhythms resynchronise to the altered host feeding rhythms when food availability is shifted, which is not mediated through rhythms in the host immune system. Our observations suggest that parasites actively control their developmental rhythms. Finally, counter to expectation, the severity of disease symptoms expressed by hosts was not affected by desynchronisation of their central and peripheral rhythms. Our study at the intersection of disease ecology and chronobiology opens up a new arena for studying host-parasite-vector coevolution and has broad implications for applied bioscience.

**Author summary:** How cycles of asexual replication by malaria parasites are coordinated to occur in synchrony with the circadian rhythms of the host is a long-standing mystery. We reveal that rhythms associated with the time-of-day that hosts feed are responsible for the timing of rhythms in parasite development. Specifically, we altered host feeding time to phase-shift peripheral rhythms, whilst leaving rhythms driven by the central circadian oscillator in the brain unchanged. We found that parasite developmental rhythms remained synchronous but changed their phase, by 12 hours, to follow the timing of host feeding. Furthermore, our results suggest that parasites themselves schedule rhythms in their replication to coordinate with rhythms in glucose in the host’s blood, rather than have rhythms imposed upon them by, for example, host immune responses. Our findings reveal a novel relationship between hosts and parasites that if disrupted, could reduce both the severity and transmission of malaria infection.

## Introduction

The discovery of daily rhythms in parasites dates back to the Hippocratic era and a taxonomically diverse range of parasites (including fungi, helminths, Coccidia, nematodes, trypanosomes, and malaria parasites [1-6]) display rhythms in development and several behaviours. Yet, how rhythms in many parasite traits are established and maintained remains mysterious, despite their significance, as these traits underpin the replication and transmission of parasites [7]. For example, metabolic rhythms of *Trypanosoma brucei* have recently been demonstrated to be under the control of an oscillator belonging to the parasite, but the constituents of this oscillator are unknown [8]. In most organisms, endogenous circadian oscillators (“clocks”) involve transcription-translation feedback loops whose timing is synchronised to external cues, such as light-dark and feeding-fasting cycles [9,10] but there is generally little homology across taxa in the genes underpinning oscillators. Multiple, convergent, evolutionary origins for circadian oscillators is thought to be explained by the fitness advantages of being able to anticipate and exploit predictable daily changes in the external environment, as well as keeping internal processes optimally timed [11,12]. Indeed, the 2017 Nobel Prize in Physiology/Medicine recognises the importance of circadian oscillators [13,14].

The environment that an endoparasite experiences inside its host is generated by many rhythmic processes, including daily fluctuations in the availability of resources, and the nature and strength of immune responses [15,16]. Coordinating development and behaviour with rhythms in the host (or vector) matters for parasite fitness [17]. For example, disrupting synchrony between rhythms in the host and rhythms in the development of malaria parasites during asexual replication reduces parasite proliferation and transmission potential [18,19]. Malaria parasites develop synchronously during cycles of asexual replication in the host’s blood and each developmental stage occurs at a particular time-of-day. The synchronous bursting of parasites at the end of their asexual cycle, when they release their progeny to infect new red blood cells, causes fever with sufficient regularity (24, 48, or 72 hourly, depending on the species) to have been used as a diagnostic tool. Malaria parasites are assumed to be intrinsically arrhythmic and mathematical modelling suggests that rhythms in host immune effectors, particularly inflammatory responses, could generate rhythms in the development of malaria parasites via time-of-day-specific killing of different parasite developmental stages [20,21]. However, the relevant processes operating within real infections remain unknown [22].

Our main aim is to use the rodent malaria parasite *Plasmodium chabaudi* to ask which circadian rhythms of the host are involved in scheduling rhythms in parasite development. In the blood, *P. chabaudi* develops synchronously and asexual cycles last 24 hours, bursting to release progeny (schizogony) in the middle of the night when mice are awake and active. We perturbed host feeding time (timing of food intake), which is known to desynchronise the phase of rhythms from the host’s central and peripheral oscillators, and we then examined the consequences for parasite rhythms. In mammals, the central oscillator in the brain (suprachiasmatic nuclei of the hypothalamus, SCN), is entrained by light [10,23]. The SCN is thought to shape rhythms in physiology and behaviour (peripheral rhythms) by entraining peripheral oscillators via hormones such as glucocorticoids [24]. However, oscillators in peripheral tissues are self-sustained and can also be entrained by several non-photic cues, such as the time-of-day at which feeding occurs [25,26]. Thus, eating at the wrong time-of-day (e.g. diurnal feeding in nocturnal mice) leads to altered timing of oscillators, and their associated rhythms in peripheral tissues. This phase-shift is particularly apparent in the liver where an inversion in the peak phase of expression of the circadian oscillator genes *Per1* and *Per2* occurs [26]. Importantly, eating at the wrong time-of-day does not alter rhythmic outputs from the central oscillator [25].

In murine hosts with an altered (diurnal) feeding schedule, the development rhythms of parasites remained synchronous but became inverted relative to the rhythms of parasites in hosts fed at night. Thus, feeding-related outputs from the hosts peripheral timing system, not the SCN, are responsible for the timing (phase) of parasite rhythms. We also reveal that the inversion of parasite rhythms corresponds to a phase-shift in blood glucose rhythms. That parasites remain synchronous during the rescheduling of their rhythm coupled with evidence that immune responses do not set the timing of parasite rhythms, suggests parasites are responsible for scheduling their developmental rhythm, and may express their own circadian rhythms and/or oscillators. Furthermore, our perturbed feeding regimes are comparable to shift work in humans. This lifestyle is well-known for increasing the risk of non-communicable diseases (cancer, type 2 diabetes etc. [27]) but our data suggest the severity of malaria infection (weight loss, anaemia) is not exacerbated by short-term desynchronisation of the central and peripheral oscillators.

## Results & Discussion

First, we examined the effects of changing the time of food intake on the phasing of circadian rhythms in host body temperature and locomotor activity (Fig 1). Body temperature is a commonly used phase marker of circadian timing because core body temperature increases during activity and decreases during sleep [28,29]. Mice were given access to food for 12 hours in each circadian cycle, either in the day (LF, light fed) or night (DF, dark fed). All food was available *ad libitum* and available from ZT 0-12 (ZT refers to ‘Zeitgeber Time’; ZT 0 is the time in hours since lights on) for LF mice, and from ZT 12-24 for DF mice. All experimental mice were entrained to the same reversed photoperiod, lights on: 7pm (ZT 0/24), lights off: 7am (ZT 12), for 2 weeks prior to starting the experiment (Fig 1).

**Fig 1.**
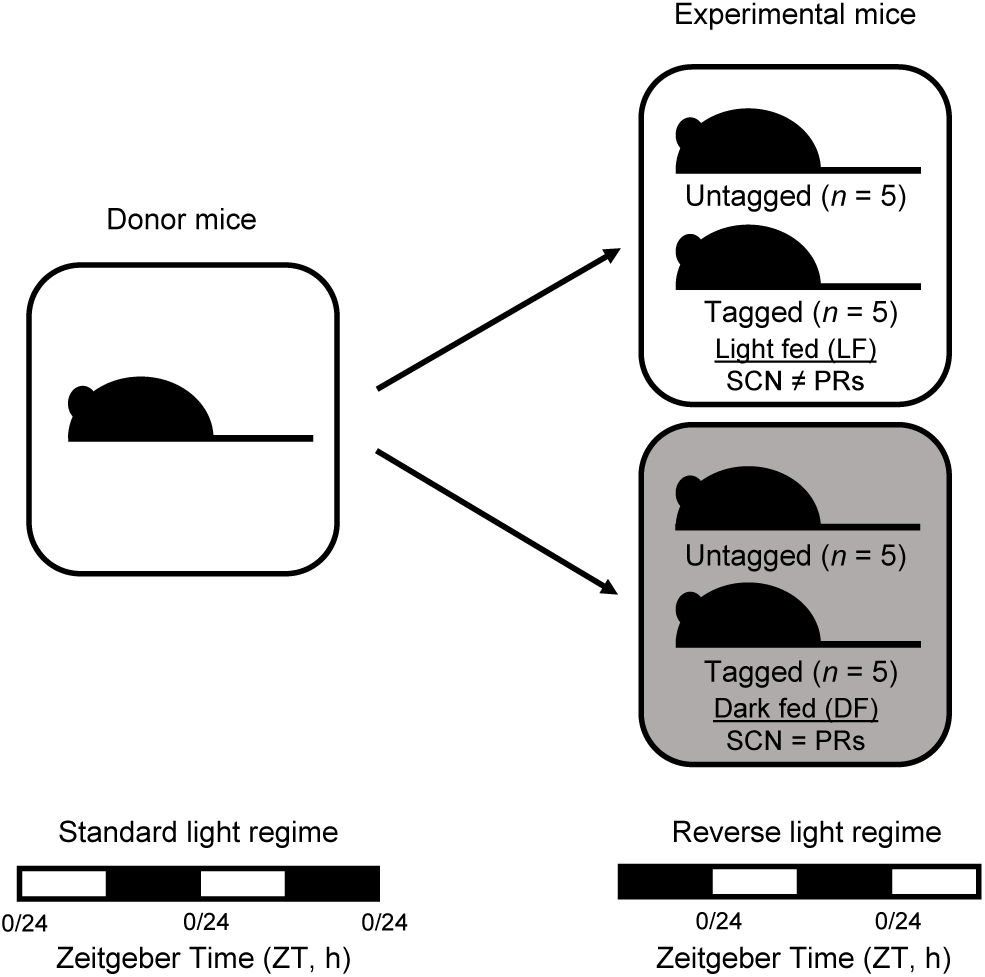
Experimental design, feeding time. Infections were initiated with parasites raised in donor mice entrained to a standard light regime [lights on: 7am (ZT 0/24) and lights off: 7pm (ZT 12)] and used to create experimental infections in hosts entrained to a reverse light regime of 12-hours light: 12-hours dark [lights on: 7pm (ZT 0/24), lights off: 7am (ZT 12); ZT is Zeitgeber Time: hours after lights on], leading to a 12-hour phase difference in SCN rhythms of donor and host, and subsequently, parasite infections (see Materials and Methods for the rationale). Hosts were then assigned to one of the two treatment groups. One group (N=10) were allowed access to food between ZT0 and ZT12 (“light fed mice”, LF, food access during the day) and the other group (N=10) allowed access to food between ZT12 and ZT0 (“dark fed mice”, DF, food access during the night). Body temperature and locomotor activity were recorded from a subset of RFID “tagged” mice in each group (N=5 per group). Changing feeding time (day time feeding of nocturnal mice) desynchronises rhythmic outputs from the central (SCN) oscillator and the peripheral (peripheral rhythms, PRs) oscillators (“SCN ≠ PRs”), whereas the SCN and peripheral rhythms remain synchronised in mice fed at night (“SCN = PRs”).

We found a significant interaction between feeding treatment (LF or DF) and the time-of-day (day (ZT 0-12) or night (ZT 12-24)) that mice experience elevated body temperatures (χ^2^_(5,6)_ = 75.89, *p* < 0.0001) and increase their locomotor activity (χ^2^_(5,6)_ = 39.57, *p* < 0.0001; S1 Table). Specifically,DF mice have elevated body temperature and are mostly active during the night (as expected) whereas LF mice show no such day-night difference in body temperature and locomotor activity, due to a lack of night time elevation in both measures where food and light associated activity are desynchronised (Fig 2). We also find the centres of gravity (CoG; a general phase marker of circadian rhythms, estimated with CircWave), are slightly but significantly earlier in LF mice for both body temperature (approximately 2 hours advanced: χ^2^_(3,4)_ = 28.17, *p* < 0.0001) and locomotor activity (approximately 4 hours advanced: χ^2^_(3,4)_ = 27.32, *p* < 0.0001) (S1 Table). Therefore, the LF mice experienced a significant change in the daily profile of activity, which is reflected in some phase advance (but not inversion) relative to DF mice, and significant disruption to their body temperature and locomotor activity rhythms, particularly during the night. Because an altered feeding schedule does not affect the phase of the SCN [25], our data suggest that rhythms in body temperature and locomotor activity in LF mice are shaped by both rhythms in feeding and the light-dark cycle [30]. Finally, the body weight of LF and DF mice did not differ significantly after 4 weeks (χ^2^_(3,4)_ = 0.02, *p* = 0.9) and both groups equally gained weight during the experiment (S1 Fig), corroborating that LF mice were not calorie restricted.

**Fig 2.**
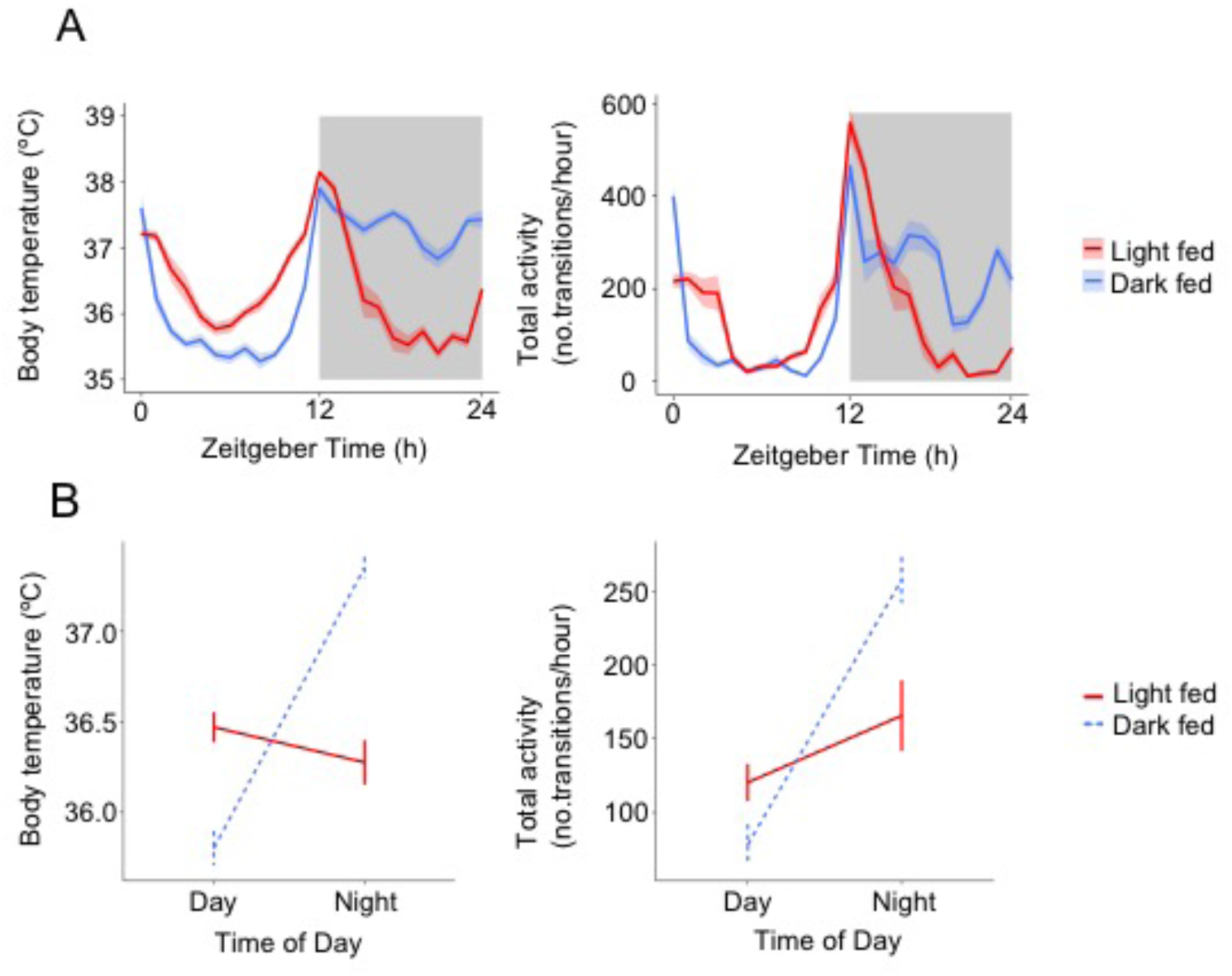
Feeding nocturnal mice in the day time disrupts rhythms in body temperature and locomotor activity. (A) Hourly mean ± SEM body temperature and locomotor activity (number of transitions per hour is the average number of movements a mouse makes in an hour, between antennae on the Home Cage Analysis system, see Materials and Methods) and (B) interaction between time-of-day and treatment group on body temperature and locomotor activity (calculating the mean temperature/activity across the day, ZT 0-12, and night, ZT 12-24, ± SEM) averaged from 48 hours of monitoring mice before infection. N=5 for each of the light fed (LF, red) and dark fed (DF, blue) groups. Light and dark bars indicate lights on and lights off (lights on: ZT 0/24, lights off: ZT 12).

Having generated hosts in which the phase relationship between the light-entrained SCN and food-entrained rhythms are altered (LF mice) or not (DF mice), we then infected all mice with the rodent malaria parasite *Plasmodium chabaudi adami* genotype DK (Fig 1) from donor mice experiencing a light-dark cycle 12 hours out of phase with the experimental host mice. After allowing the parasite’s developmental rhythms to become established (see Materials and Methods) we compared the rhythms of parasites in LF and DF mice. We hypothesised that if parasite rhythms are solely determined by rhythms driven by the host’s SCN (which are inverted in the host mice compared to the donor mice), parasite rhythms would equally shift and match in LF and DF mice because both groups of hosts were entrained to the same light-dark conditions. Yet, if rhythms in body temperature or locomotor activity directly or indirectly (via entraining other oscillators) contribute to parasite rhythms, we expected that parasite rhythms would differ between LF and DF hosts. Further, if feeding directly or indirectly (via food-entrained oscillators) drives parasite rhythms, we predicted that parasite rhythms would become inverted (Fig 1).

In the blood, *P. chabaudi* parasites transition through five developmental stages during each (~24hr) cycle of asexual replication (Fig 3A) [6,31]. We find that four of the five developmental stages (rings, and early-, mid-, and late-trophozoites) display 24hr rhythms in both LF and DF mice (Fig 3B, S2 Table, S2 Fig). The fifth stage - schizonts - appear arrhythmic but this stage sequesters in the host’s tissues [32,33] and so, are rarely collected in venous blood samples. Given that all other stages are rhythmic, and that rhythms in ring stages likely require their parental schizonts to have been rhythmic, we expect schizonts are rhythmic but that sequestration prevents a reliable assessment of their rhythms.

**Fig 3.**
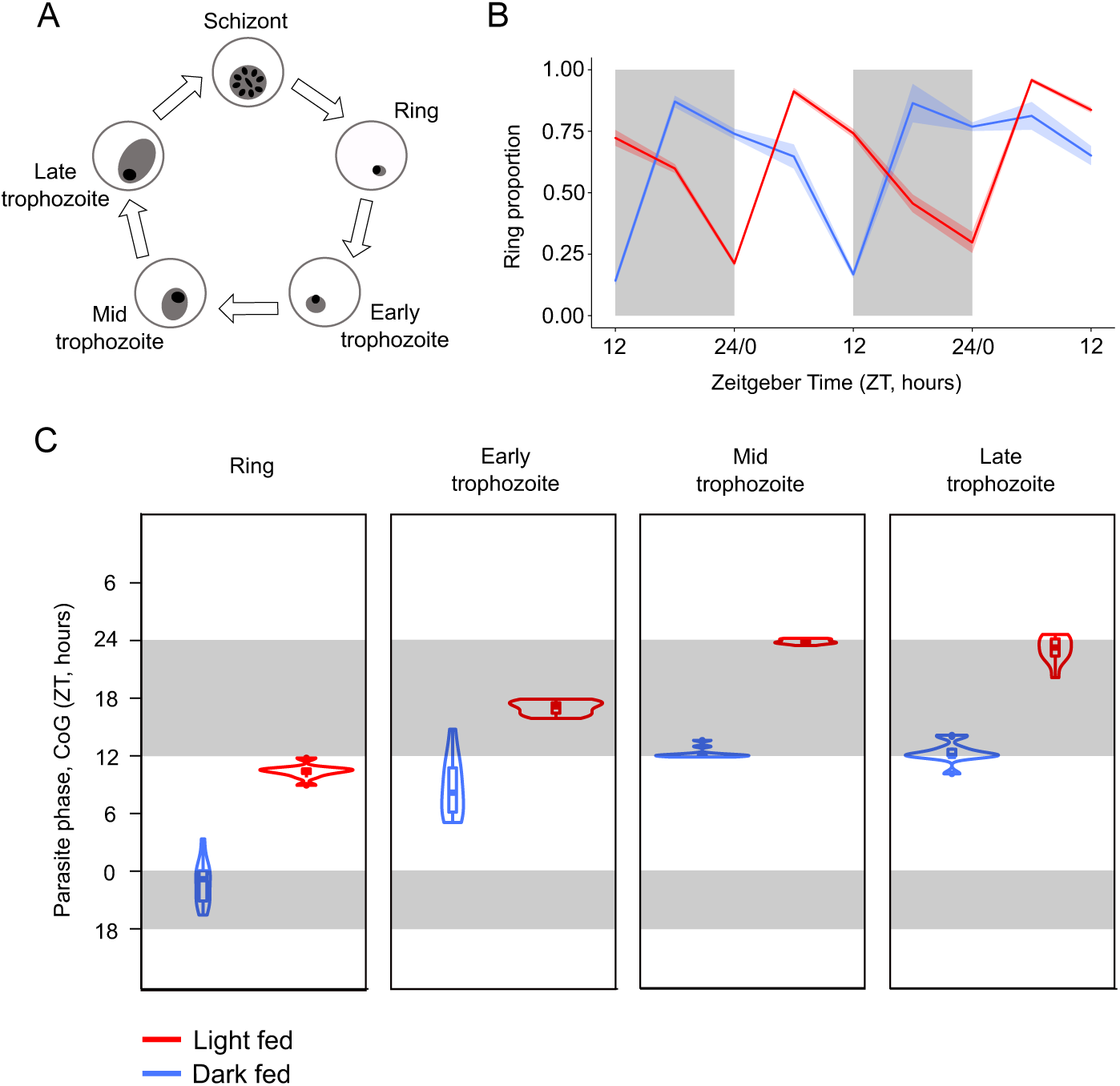
Parasite rhythms are inverted in hosts fed during the day compared to the night. (A) The asexual cycle of malaria parasites is characterised by five morphologically distinct developmental stages (ring, early trophozoite, mid trophozoite, late trophozoite) differentiated by parasite size within the red blood cell, the size and number of nuclei, and the appearance of haemozoin [31]. (B) Mean ± SEM (N=10 per group) proportion of observed parasites in the blood at ring stage in light fed mice (red; allowed access to food during the day, between ZT 0 and ZT 12) and dark fed mice (blue; allowed access to food during the night, between ZT 12 and ZT 24). The proportion of parasites at ring stage in the peripheral blood is highest at night (ZT 22) in dark fed mice but in the day (ZT 10) for light fed mice, illustrating the patterns observed for all other (rhythmic) stages (see Fig S2). (C) CoG (estimate of phase) in ZT (h) for each rhythmic parasite stage in the blood. Each violin illustrates the median ± IQR overlaid with probability density (N=10 per group). The height of the violin illustrates the variation in the timing of the CoG between mice and the width illustrates the frequency of the CoGs at particular times within the distribution. Sampling occurred every 6 hours days 6-8 post infection. Light and dark bars indicate lights on and lights off (lights on: ZT 0, lights off: ZT 12).

The CoG estimates for ring, and early-, mid-, and late-trophozoite stages are approximately 10-12 hours out-of-phase between the LF and DF mice (Fig 3B,C, S2 Table). For example, rings peak at approximately ZT 10 in LF mice and peak close to ZT 23 in DF mice. The other stages peak in sequence. Schizogony (when parasites burst to release their progeny) occurs immediately prior to reinvasion, therefore we expect it occurs during the day for the LF mice and night for DF mice [7]. The almost complete inversion in parasite rhythms between LF and DF mice demonstrates that feeding-related rhythms are responsible for the phase of parasite rhythms, with little to no apparent contribution from the SCN and/or the light: dark cycle.

Changing the feeding time of nocturnal mice to the day time has similarities with shift work in diurnal humans [34]. This lifestyle is associated with an increased risk of acquiring non-communicable diseases (e.g. cancer, diabetes) [35] and has been recapitulated in mouse models [e.g. 36,37,38]. In contrast, in response to perturbation of their feeding rhythm, infections are not more severe in hosts whose circadian rhythms are desynchronised (i.e. LF hosts). Specifically, all mice survived infection and virulence (measured as host anaemia; reduction in red blood cells) of LF and DF infections is not significantly different (comparing minimum red blood cell density, χ^2^_(3,4)_ = 0.11, *p* = 0.74; S3A Fig). As described above, changes in body mass were not significantly different between treatments (S1 Fig). Using a longer-term model for shift work may reveal differences in infection severity, especially when combined with the development of non-communicable disease.

There are no significant differences between parasite densities in LF and DF hosts during infections (LF versus DF on day 6 post infection, χ^2^_(3,5)_ = 0.66, *p* = 0.42, S3B Fig). This can be explained by both groups being mismatched to the SCN of the host, which we have previously demonstrated to have negative consequences for *P. chabaudi* [18]. Our previous work was carried out using *P. chabaudi* genotype AJ so is not directly comparable to our results presented here, because DK is a less virulent genotype [39]. Instead, a comparison of our results to data collected previously for genotype DK, in an experiment where SCN rhythms of donor and host mice were matched (see Materials and Methods; infections were initiated with the same strain, sex, and age of mice, the same dose at ring stage) reveals a cost of mismatch of donor and host entrainment. Specifically, parasite density on day 6 (when infections have established but before parasites start being cleared by host immunity) is significantly lower in infections mismatched to the SCN (LF and DF) compared to infections matched to the SCN (χ^2^_(3,5)_ = 16.71, *p* = 0.0002, difference = 2.21e+10 parasites per ml blood) (see S4A Fig). In keeping with a difference in parasite replication, hosts with matched infections reach lower red blood cell densities (χ^2^_(3,5)_ = 18.87, *p* < 0.0001, mean difference = 5.29e+08 red blood cells per ml blood).

The mismatched and matched infections compared above also differ in whether hosts had food available throughout the 24-hour cycle or for 12 hours only (LF and DF). Restricting food to 12 hours per day does not affect host weight (S1 Fig) and mice still undergo their main activity bout at lights off even when food is available all the time. Therefore, we propose that rather than feeding duration, mismatch to the host SCN for as few as 5 cycles is costly to parasite replication and reduces infection severity. Because peripheral and SCN driven rhythms are usually in synchrony, we suggest parasites use information from food-entrained oscillators, or metabolic processes, to ensure their development is timed to match the host’s SCN rhythms.

Instead of organising their own rhythms (i.e. using an “oscillator” whose time is set by a “Zeitgeber” or by responding directly to time-of-day cues), parasites may allow outputs of food-entrained host oscillators to enforce developmental rhythms. Previous studies have focused on rhythmic immune responses as the key mechanism that schedules parasite rhythms (via developmental-stage and time-of-day specific killing [20,21]). Evidence that immune responses are rhythmic in naïve as well as infected hosts is increasing [15,16], but the extent to which peripheral/food-entrained oscillators and the SCN drive immune rhythms is unclear. Nonetheless, we argue that rhythms in host immune responses do not play a significant role in scheduling parasites for the following reasons: First, mismatch to the host’s peripheral rhythms (which occurs in DF mice but not LF mice as a feature of our experimental design) does not cause a significant reduction in parasite number (S3B Fig), demonstrating that stage-specific killing cannot cause the differently phased parasite rhythms in LF and DF mice. Second, while changing feeding time appears to disrupt some rodent immune responses [40,41], effectors important in malaria infection, including leukocytes in the blood, do not entrain to feeding rhythms [42,43]. Third, inflammatory responses important for killing malaria parasites are upregulated within hours of blood stage infection [44] so their footprint on parasite rhythms should be apparent from the first cycles of replication [19]. In contrast, rhythms of parasites in LF and DF mice do not significantly diverge until 5-6 days post infection, after 5 replication cycles (S3 Table, Fig 4). Fourth, an additional experiment (see Materials and Methods) reveals that rhythms in the major inflammatory cytokines that mediate malaria infection (e.g. IFN-gamma and TNF-alpha: [45,46,47,48]) follow the phase of parasite rhythms (Fig 5), with other cytokines/chemokines also experiencing this phenomenon (S5 Fig). Specifically, mice infected with *P. chabaudi* genotype AS undergoing schizogony at around midnight (ZT17), produce peaks in the cytokines IFN-gamma and TNF-alpha at ZT21 and ZT19 respectively. Whereas mice infected with mismatched parasites undergoing schizogony around ZT23 (6 hours later), experience 3-6 hour delays in the peaks of IFN-gamma and TNF-alpha (IFN-gamma: ZT0, TNF-alpha: ZT1). Thus, even if parasites at different development stages differ in their sensitivity to these cytokines, these immune rhythms could only serve to increase synchrony in the parasite rhythm but not change its timing.

**Fig 4.**
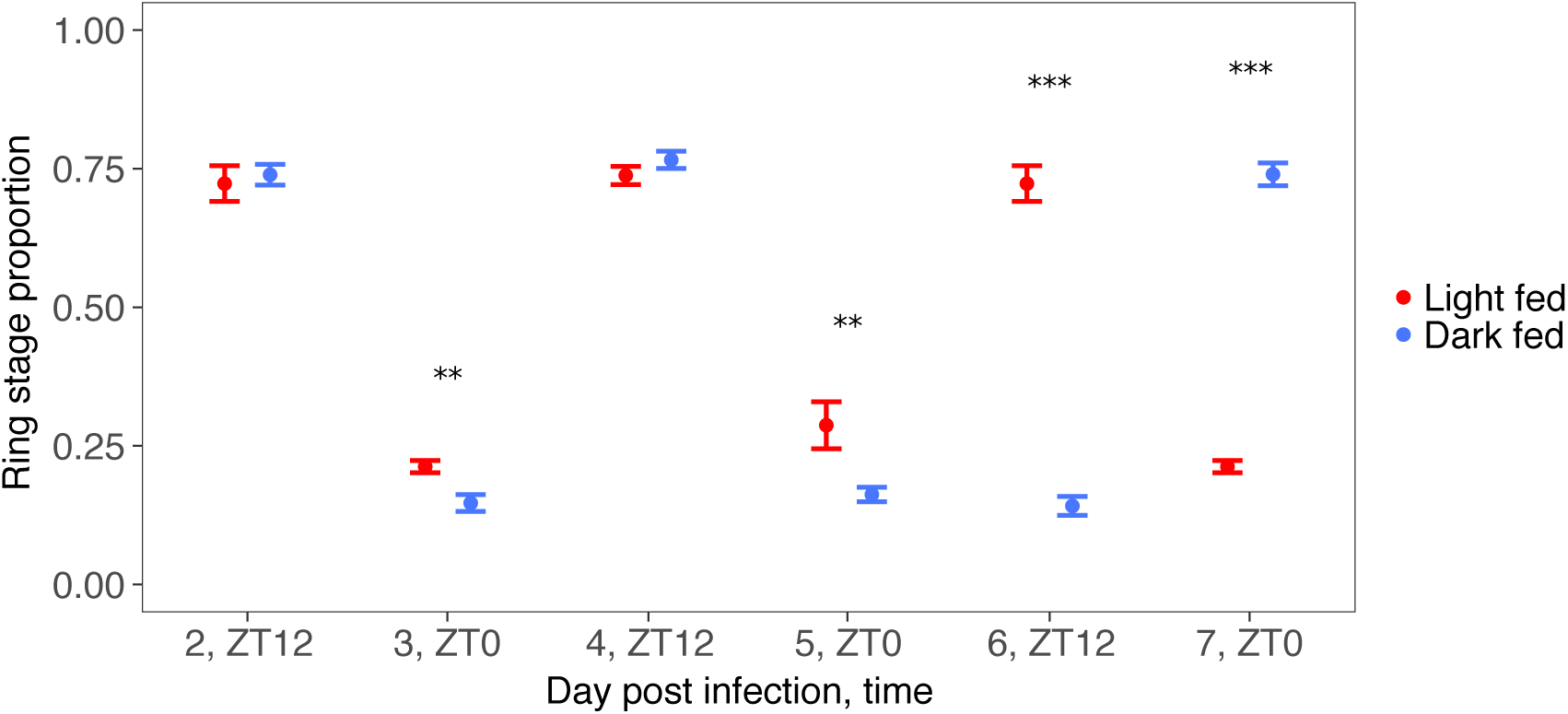
Parasite rhythms in light and dark fed mice significantly diverge by day 5-6 post infection. The proportion of ring stage parasites across infections (light fed mice, red, and dark fed mice, blue) as a phase marker reveals that rhythms of parasites in light fed mice (red) and dark fed mice (blue) diverge. Mice were sampled at ZT 12 on days 2, 4 and 6 and at ZT 0 on days 3, 5 and 7 post infection (see Fig 3 and S2 Fig). Consistent significant differences (**, *p* < 0.05; ***, *p* < 0.001) between feeding treatments begins on day 5. By days 6-7 post infection, rings in light fed mice are present at ZT12 while rings in dark fed mice are present at ZT 0, indicating that parasites in dark fed mice have rescheduled. Ring stages are presented as the phase marker because this is the most accurately quantified stage but other stages follow a similar pattern (S3 Table). Mean ± SEM is plotted and N=10 for each treatment group.

**Fig 5.**
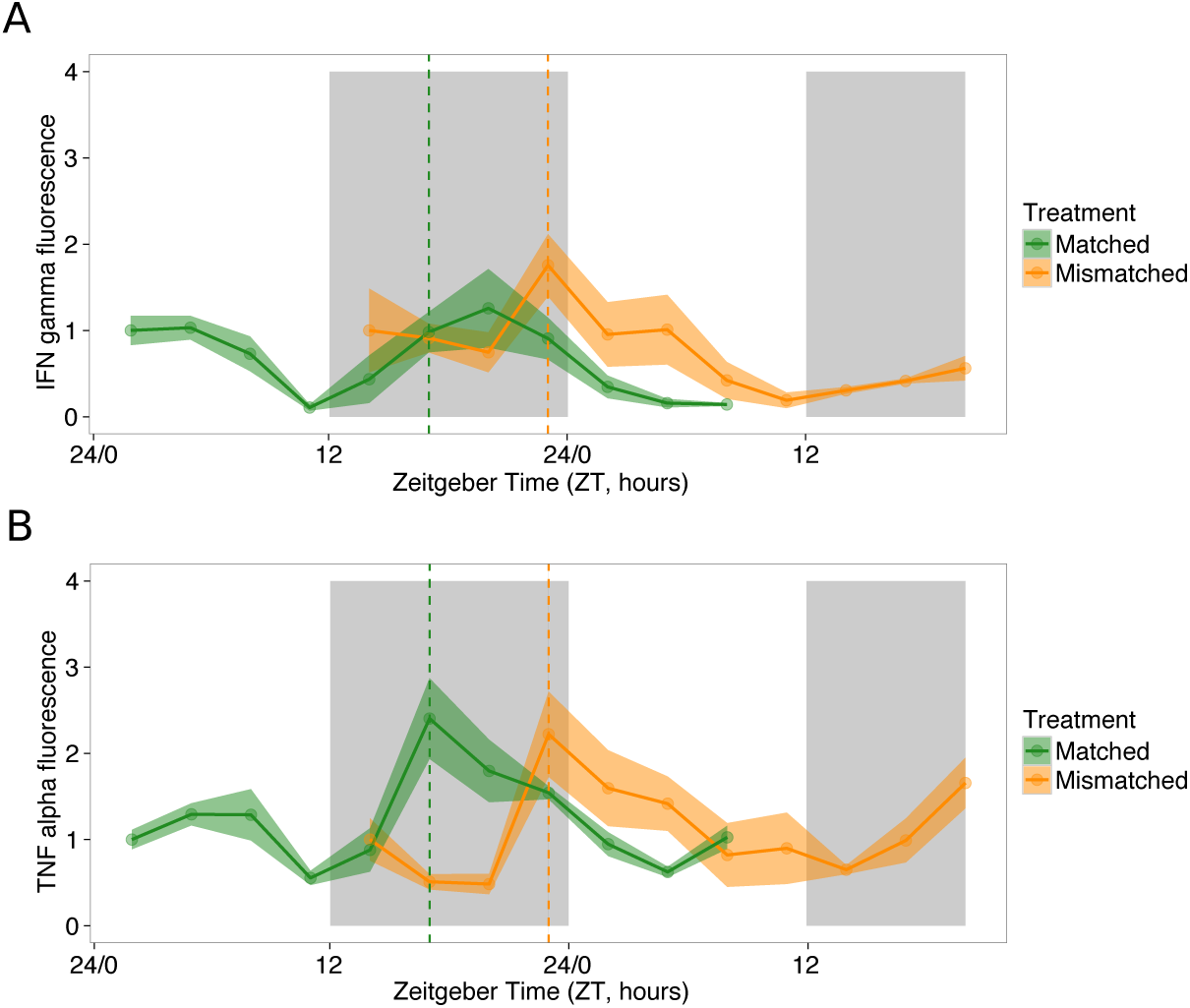
Rhythms in inflammatory cytokines follow rhythms in parasite development. Mean ± SEM (N=4 per time point) for cytokines (A) IFN-gamma and (B) TNF-alpha for parasites matched and mismatched to the SCN rhythms of the host (matched: green, mismatched: orange). Sampling occurred every 3 hours on days 4-5 post infection. Matched parasites undergo schizogony around ZT 17, (indicated by green dashed line) and mismatched parasites undergo schizogony 6 hours later, around ZT 23 (indicated by orange dashed line). IFN-gamma peaks at ZT 21.29 in matched infections (green) and at ZT 0 in mismatched infections (orange). TNF-alpha peaks at ZT 19.26 in matched infections (green) and at ZT 1.29 in mismatched infections (orange). Light and dark bars indicate lights on and lights off (lights on: ZT 0, lights off: ZT 12).

More in-depth analysis of LF and DF infections provides further support that parasites actively organise their developmental rhythms. We examined whether parasites in DF mice maintain synchrony and duration of different developmental stages during rescheduling to the host’s SCN rhythms. Desynchronisation of oscillators manifests as a reduction in amplitude in rhythms that are driven by more than one oscillator (e.g. parasite and host oscillator). No loss in amplitude suggests that parasites shift their timing as a cohort without losing synchrony. Parasite rhythms in LF and DF mice did not differ significantly in amplitude (χ^2^_(6,7)_ = 1.53, *p* = 0.22, S4A Table) and CoGs for sequential stages are equally spaced (χ^2^_(10,18)_ = 11.75, *p* = 0.16, S2 Table) demonstrating that parasite stages develop at similar rates in both groups. The rhythms of parasites in LF and DF mice were not intensively sampled until days 6-8 PI, raising the possibility that parasites lost and regained synchrony before this. Previously collected data for *P. chabaudi* genotype AS infections mismatched to the host SCN by 12 hours that have achieved a 6-hour shift by day 4 PI also exhibit synchronous development (S4B Table and S6 Fig), suggesting that parasites reschedule in synch.

That parasite rhythms do not differ significantly between LF and DF mice until day 5-6 post infection (Fig 4) could be explained by the parasites experiencing a phenomenon akin to jet lag. Jet lag results from the fundamental, tissue-specific robustness of circadian oscillators to perturbation, which slows down the phase shift of individual oscillators to match a change in ‘time-zone’ [10]. We propose that the most likely explanation for the data gathered from our main experiment for genotype DK, and that collected previously for AJ and AS, is that parasites possess intrinsic oscillators that shift collectively, in a synchronous manner, by a few hours each day, until they re-entrain to the new ‘time-zone’. Because there is no loss of amplitude of parasite rhythms, it is less likely that individual parasites possess intrinsic oscillators that re-entrain at different rates to the new ‘time-zone’. The recently demonstrated ability of parasites to communicate decisions about asexual to sexual developmental switches [49] could also be involved in organising asexual development.

If parasites have evolved a mechanism to keep time and schedule their rhythms, what external information might they synchronise to? Despite melatonin peaks in lab mice being brief and of low concentration [50,51], the host’s pineal melatonin rhythms have been suggested as a parasite time cue [52]. However, we can likely rule pineal melatonin, and other glucocorticoids, out because they are largely driven by rhythms of the SCN, which follow the light-dark cycle and have not been shown to phase shift by 12 hours as a result of perturbing feeding timing [25]; some glucocorticoid rhythms appear resistant to changing feeding time [53]. Whether extra-pineal melatonin, produced by the gut for example [54], could influence the rhythms of parasites residing in the blood merits further investigation. Body temperature rhythms have recently been demonstrated as a Zeitgeber for an endogenous oscillator in trypanosomes [8]. Malaria parasites are able to detect and respond to changes in environmental temperature to make developmental transitions in the mosquito phase of their lifecycle [55,56], and may deploy the same mechanisms to organise developmental transitions in the host. Body temperature rhythms did not fully invert in LF mice but they did exhibit unusually low (i.e. day time) temperatures at night. Thus, for body temperature to be a time-of-day cue or Zeitgeber it requires that parasites at early developmental stages (e.g. rings or early trophozoites) are responsible for time-keeping because they normally experience low temperatures during the day when the host is resting. The same logic applies to rhythms in locomotor activity because it is very tightly correlated to body temperature (Pearson’s correlation R=0.85, 95% CI: 0.82-0.88). Locomotor activity affects other rhythms, such as physiological oxygen levels (daily rhythms in blood and tissue oxygen levels), which can reset circadian oscillators [57] and have been suggested as a time cue for filarial nematodes [4].

Feeding rhythms were inverted in LF and DF mice and so, the most parsimonious explanation is that parasites are sensitive to rhythms related to host metabolism and/or food-entrained oscillators. Malaria parasites have the capacity to actively alter their replication rate in response to changes in host nutritional status [58]. Thus, we propose that parasites also possess a mechanism to coordinate their development with rhythms in the availability of nutritional resources in the blood. Rhythms in blood glucose are a well-documented consequence of rhythms in feeding timing [59] and glucose is an important resource for parasites [60]. We performed an additional experiment to quantify blood glucose rhythms in (uninfected) LF and DF mice (Fig 6A,B). Despite the homeostatic regulation of blood glucose, we find its concentration varies across the circadian cycle, and is borderline significantly rhythmic in DF mice (*p* = 0.07, peak time = ZT17.84, estimated with CircWave) and follows a significantly 24-hour pattern in LF mice (*p* < 0.0001, peak time = ZT8.78). Glucose rhythms/patterns are shaped by feeding regime (time-of-day: feeding treatment χ^2^_(18,32)_ = 45.49, *p* < 0. 0001). Specifically, during the night, DF mice have significantly higher blood glucose than LF mice (t = 3.41, *p* = 0.01, difference 20.6mg/dl±7.32) and there is a trend for LF mice to have higher blood glucose than DF mice during the day (t = −0.94, *p* = 0.78, difference 7.9mg/dl±9.86)

**Fig 6.**
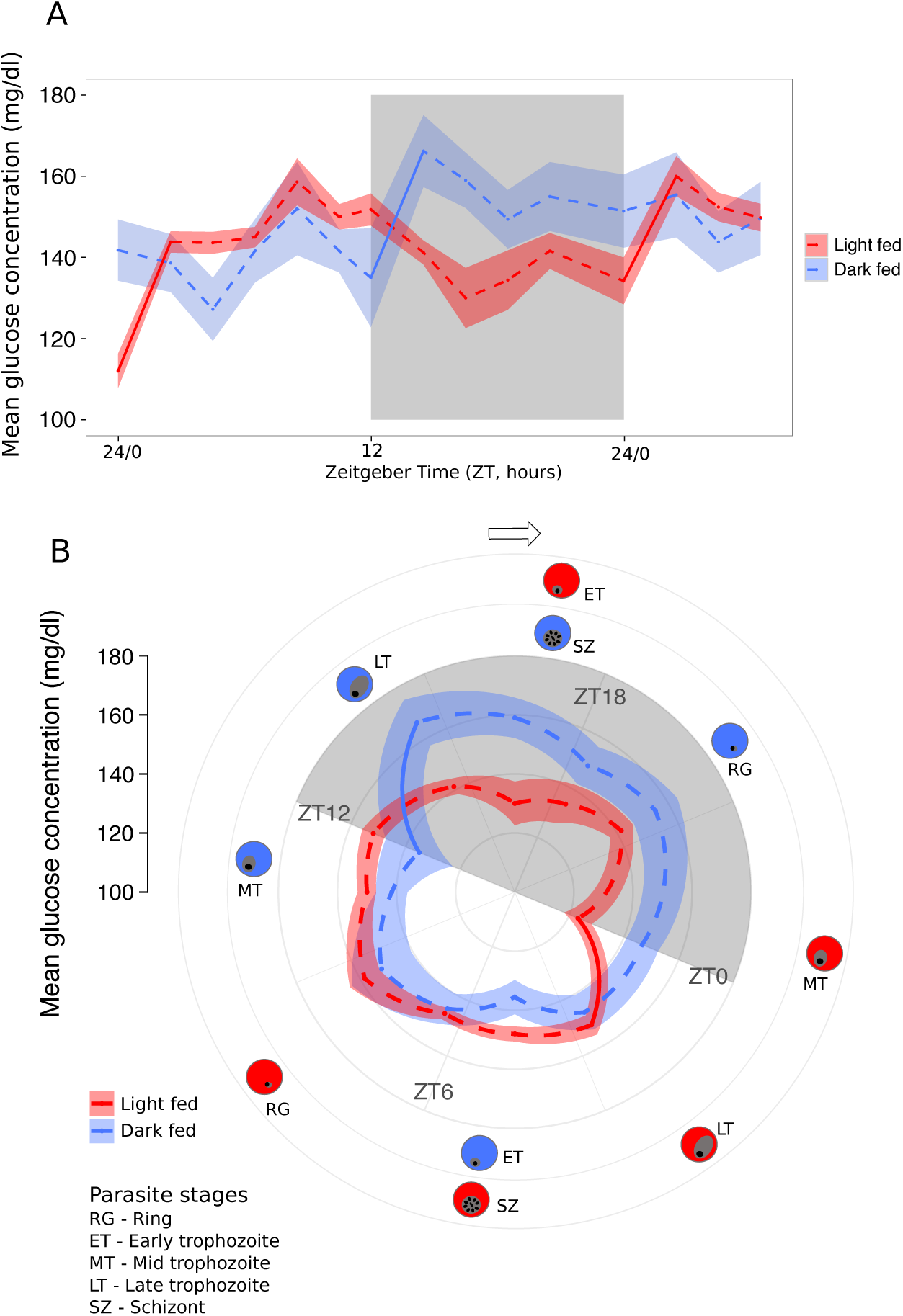
Feeding mice in the day time affects blood glucose regulation. A) Mean ± SEM (N=5 per group) for light fed mice (LF, white bars; allowed access to food from ZT 0-ZT 12) and dark fed mice (DF, grey bars; allowed access to food from ZT 12-ZT 0). Blood glucose concentration was measured every ~2 hours for 30 hours from ZT 0. Steep increases in blood glucose concentration occur as a result of the main bout of feeding in each group (i.e. just after lights on in LF mice and lights off in DF mice, illustrated by the regions with solid lines connecting before and after the main bout, see S5 Table), and suggests glucose concentration is inverted during the night. Light and dark bars indicate lights on and lights off (lights on: ZT 0, lights off: ZT 12). B) as for A, but plotted as a polar graph with corresponding developmental stages for each treatment group (red, LF; blue DF) on the perimeter.

Titrating whether glucose availability is high or low would only provide parasites with information on whether it is likely to be day or night, and a 12-hour window in which to make developmental transitions should erode synchrony, especially as glucose rhythms are weak in DF mice. Instead, parasites may use the sharp rise in blood glucose that occurs in both LF and DF mice after their main bout of feeding as a cue for dusk (S5 Table; regions with solid lines connecting before and after feeding in Fig 6). In line with the effects of feeding timing we observe in mice, a recent study of humans reveals that changing feeding time can induce a phase-shift in glucose rhythms, but not insulin rhythms [43]. Alternatively, parasites may be sensitive to fluctuations in other factors due to rhythms in food intake, such as amino acids [61] or other rhythmic metabolites that appear briefly in the blood after feeding, changes in oxygen consumption, blood pressure or blood pH [62,63].

In summary, we show that peripheral, food-entrained host rhythms, but not central, light-entrained host rhythms are responsible for the timing of developmental transitions during the asexual replication cycles of malaria parasites. Taken together, our observations suggest that parasites have evolved a time-keeping mechanism that uses daily fluctuations in resource availability (e.g. glucose) as a time-of-day cue or Zeitgeber to match the phase of asexual development to the host’s SCN rhythms. Why coordination with the SCN is important remains mysterious. Uncovering how parasites tell the time could enable an intervention (ecological trap) to “trick” parasites into adopting suboptimal rhythms for their fitness.

## Materials and Methods

We conducted an experiment to investigate whether host peripheral rhythms or those driven by the SCN affect rhythms in the asexual development of malaria parasites. Our findings stimulated the analysis of four further data sets stemming from three independent experiments. Here, we detail the approach used for our main experiment “Effect of feeding time on parasite rhythms” before briefly outlining the approaches used in the analyses of additional data “Costs of mismatch to host SCN rhythms”, “Rhythms in cytokines during malaria infection”, “Synchrony during rescheduling” and “Effect of feeding time on blood glucose rhythms”.

### Effect of feeding time on parasite rhythms

#### Experimental design

Both LF (“light-fed mice”, access to food during the day, ZT 0-12) and DF (“dark-fed mice”, access to food during the night, ZT 12-0) mice were kept in the same light-dark cycle to ensure the phase of their central oscillators did not differ (because the SCN is primarily entrained by light [23]) (Fig 1). Changing host feeding time in LF mice created an in-host environment where peripheral rhythms associated with feeding are out of phase with the SCN, but in phase in DF mice. Every 12 hours, food was added/removed from cages and the cages thoroughly checked for evidence of hoarding, which was never observed. All experimental infections were initiated with parasites from donor mice in light-dark cycles that were out of phase with the experimental host’s light-dark cycles by 12 hours, leading to a 12-hour phase difference in SCN entrainment of donor and host. Specifically, infections were initiated with ring stage parasites (which appear in the early morning) collected from donor mice and injected immediately into host mice which experiencing their evening. Parasites that are mismatched by 12 hours to mice with synchronised SCN and peripheral rhythms (i.e. DF mice) take around one week to reschedule [64,65,18]. Therefore, if peripheral rhythms but not SCN rhythms, affect parasite rhythms, by starting infections with mismatched parasites we expected that parasites in DF mice would reschedule within 7 days whereas rhythms in the LF mice would not change (or change less). Because rhythms generally return to their original state after perturbation faster than they can be shifted from homeostasis [66], studying the change in rhythms of mismatched parasites ensured we could observe any divergence between parasite rhythms in LF and DF mice before host immune responses and anaemia clear infections.

#### Parasites and hosts

We used 20 eight-week-old male mice, strain MF1 (in house supplier, University of Edinburgh), entrained to a reverse lighting schedule for 2 weeks before starting the experiment. After entrainment, mice were randomly allocated to one of two feeding treatments for the entire experiment (Fig 1). After 2 weeks on the assigned feeding treatment we recorded body temperature and locomotor activity for 48 hours. We used BioThermo13 RFID (radio frequency identification) tags (Biomark, Idaho, USA) in conjunction with a Home Cage Analysis system (Actual HCA, Actual Analytics Ltd, Edinburgh, Scotland), which enables body temperature and locomotor activity readings to be taken every 0.05 seconds without disturbing the animals (using a network of antennae spaced approximately 10.9 cm apart). Next, all mice were intravenously infected with 1 × 10^7^ *Plasmodium chabaudi adami* (avirulent genotype, DK) parasitised red blood cells (at ring stage). We used DK to minimise disruption to host feeding compared to infection with more virulent genotypes that cause more severe sickness [39]. All mice were blood sampled from the tail vein twice daily (ZT0 and ZT12) on days 0-5 and every 6 hours from days 6-8 post infection (PI). The densities and developmental stages of parasites in experimental infections were determined from thin blood smears (day 2 PI onwards, when parasites become visible in the blood) and red blood cell (RBC) densities by flow cytometry (Beckman Coulter).

### Costs of mismatch to host SCN rhythms

We compared the performance of parasites in our main experiment (in which infections were initiated with parasites from donor mice that were mismatched to the host’s SCN rhythms by 12 hours), to the severity of infections when infections are initiated with parasites from donor mice that are matched to the host’s SCN rhythms. Twelve infections were established in the manner used in our main experiment (eight-week-old male mice, strain MF1, intravenously infected with 1 × 10^7^ *P. chabaudi* DK parasitised RBC), except that donor SCN rhythms were matched to the experimental host’s SCN rhythm and hosts had access to food day and night. Densities of parasites were quantified from blood smears and RBC density by flow cytometry on day 6 and 9 PI, respectively. We chose to compare parasite density in matched infections to LF and DF infections on day 6 PI because parasites are approaching peak numbers in the blood (before host immunity starts to clear infections) and their high density facilitates accurate quantification when using microscopy.

### Rhythms in cytokines during malaria infection

This experiment probes whether host immune responses mounted during the early phase of malaria infection could impose development rhythms upon parasites. We entrained N=86 eight-week-old female mice, strain MF1, to either a reverse lighting schedule (lights on 7pm, lights off 7am, N=43) or a standard lighting schedule (lights on 7am, lights off 7pm, N=43). Donor mice, infected with *P. chabaudi* genotype AS, were entrained to a standard lighting schedule to generate infections matched and 12 hours mismatched relative to the SCN in the experimental mice. Mice were intravenously injected with 1 × 10^7^ parasitised RBC at ring stage. Genotype AS has intermediate virulence [39] and was used to ensure immune responses were elicited by day 4 PI. We terminally sampled 4 mice every 3 hours over 30 hours starting on day 4 PI, taking blood smears, red blood cell counts and collecting plasma for Luminex cytokine assays.

Cytokines were assayed by the Human Immune Monitoring Centre at Stanford University using mouse 38-plex kits (eBiosciences/Affymetrix) and used according to the manufacturer’s recommendations with modifications as described below. Briefly, beads were added to a 96-well plate and washed in a Biotek ELx405 washer. 60uL of plasma per sample was submitted for processing. Samples were added to the plate containing the mixed antibody-linked beads and incubated at room temperature for one hour followed by overnight incubation at 4°C with shaking. Cold and room temperature incubation steps were performed on an orbital shaker at 500-600 rpm. Following the overnight incubation, plates were washed as above and then a biotinylated detection antibody was added for 75 minutes at room temperature with shaking. Plates were washed as above and streptavidin-PE was added. After incubation for 30 minutes at room temperature a wash was performed as above and reading buffer was added to the wells. Each sample was measured as singletons. Plates were read using a Luminex 200 instrument with a lower bound of 50 beads per sample per cytokine. Custom assay control beads by Radix Biosolutions were added to each well.

### Synchrony during rescheduling

We staged the parasites from the blood smears collected from the infections used to assay cytokines (above) to investigate their synchrony during rescheduling. The infections from mismatched donor mice began 12 hours out of phase with the host SCN rhythms and the CoG for ring stage parasites reveals they had become rescheduled by 6 hours on day 4 PI. We focus on the ring stage as a phase marker – for the analysis of synchrony in these data and the divergence between LF and DF parasites – because rings are the most morphologically distinct, and so, accurately quantified, stage.

### Blood glucose concentration

In a third additional experiment, we entrained 10 eight-week-old male mice, strain MF1, to a standard lighting schedule for 2 weeks before randomly allocating them to one of two feeding treatments. One group (N=5) were allowed access to food between ZT 0 and ZT 12 (equivalent to the LF group in the main experiment) and the other group (N=5) allowed access to food between ZT 12 and ZT 0 (equivalent to the DF group). After 10 days of food restriction we recorded blood glucose concentration every 2 hours for 30 hours, using an Accu-Chek Performa glucometer.

### Data analysis

We used CircWave (version 1.4, developed by R.A. Hut; available from http://www.euclock.org/) to characterise host and parasite rhythms, and R v. 3.1.3 (The R Foundation for Statistical Computing, Vienna, Austria) for analysis of summary metrics and non-circadian dynamics of infection. Specifically, testing for rhythmicity, estimating CoG (a reference point to compare circadian rhythms) for host (body temperature, locomotor activity, blood glucose concentration) and parasite rhythms, and amplitude for parasite stage proportions, was carried out with CircWave for each individual infection. However, the cytokine data display high variation between mice (due to a single sample from each mouse) so we calculated a more robust estimate of phase than CoG by fitting a sine curve with a 24h period (using CircWave) and finding the maxima. Linear regression models and simultaneous inference of group means (using the multcomp R package) were run with R to compare summary measures that characterise rhythms, parasite performance, glucose concentration and disease severity. R was also used to construct and compared linear mixed effects models using which included mouse ID as a random effect (to account for repeated measures from each infection) to compare dynamics of parasite and RBC density throughout infections, and glucose concentration throughout the day.

### Ethics Statement

All procedures were carried out in accordance with the UK Home Office regulations (Animals Scientific Procedures Act 1986; project licence number 70/8546) and approved by the University of Edinburgh.

## Acknowledgements

We thank Philip Birget, Nicole Mideo, Petra Schneider, Bert Maier, Phil Spence and Barbara Helm for discussion, Christopher Hutton and Giles K.P. Barra for assistance, and three anonymous reviewers for their helpful and insightful comments.

